# The role of Kölliker-Fuse nucleus in breathing variability

**DOI:** 10.1101/2023.06.15.545086

**Authors:** S. John, W. Barnett, A. Abdala, D. Zoccal, J. Rubin, Y. Molkov

**Affiliations:** University of Pittsburgh, Pittsburgh, PA; Indiana University Purdue University Indianapolis, IN; University of Bristol, Bristol, UK; São Paulo State University, Araraquara, Brazil; Georgia State University, Atlanta, GA

## Abstract

The Kölliker-Fuse nucleus (KF), which is part of the parabrachial complex, participates in the generation of eupnea under resting conditions and the control of active abdominal expiration when increased ventilation is required. Moreover, dysfunctions in KF neuronal activity are believed to play a role in the emergence of respiratory abnormalities seen in Rett syndrome (RTT), a progressive neurodevelopmental disorder associated with an irregular breathing pattern and frequent apneas. Relatively little is known, however, about the intrinsic dynamics of neurons within the KF and how their synaptic connections affect breathing pattern control and contribute to breathing irregularities. In this study, we use a reduced computational model to consider several dynamical regimes of KF activity paired with different input sources to determine which combinations are compatible with known experimental observations. We further build on these findings to identify possible interactions between the KF and other components of the respiratory neural circuitry. Specifically, we present two models that both simulate eupneic as well as RTT-like breathing phenotypes. Using nullcline analysis, we identify the types of inhibitory inputs to the KF leading to RTT-like respiratory patterns and suggest possible KF local circuit organizations. When the identified properties are present, the two models also exhibit quantal acceleration of late-expiratory activity, a hallmark of active expiration featuring forced exhalation, with increasing inhibition to KF, as reported experimentally. Hence, these models instantiate plausible hypotheses about possible KF dynamics and forms of local network interactions, thus providing a general framework as well as specific predictions for future experimental testing.

**Key points:** The Kölliker-Fuse nucleus (KF), a part of the parabrachial complex, is involved in regulating normal breathing and controlling active abdominal expiration during increased ventilation. Dysfunction in KF neuronal activity is thought to contribute to respiratory abnormalities seen in Rett syndrome (RTT). This study utilizes computational modeling to explore different dynamical regimes of KF activity and their compatibility with experimental observations. By analyzing different model configurations, the study identifies inhibitory inputs to the KF that lead to RTT-like respiratory patterns and proposes potential KF local circuit organizations. Two models are presented that simulate both normal breathing and RTT-like breathing patterns. These models provide plausible hypotheses and specific predictions for future experimental investigations, offering a general framework for understanding KF dynamics and potential network interactions.

## 1 Introduction

Breathing is an automatic process produced and shaped by the respiratory central pattern generator (rCPG), which comprises several neural structures in the brainstem (Smith et al., 2007). Extensive work has characterized the synaptic and ionic mechanisms that generate and regulate the activity of ventromedullary respiratory neurons and how these mechanisms affect breathing rhythm and pattern. This study focuses on a major counterpart to the medullary rCPG: the pontine Kölliker-Fuse nucleus (KF). The KF is part of the parabrachial complex in the dorsolateral pons and is formed by a collection of neurons surrounding the superior cerebellar peduncle (Varga et al., 2020). Although it was first described as the inspiratory off-switch center, compelling evidence indicates that the KF activity critically contributes to maintaining the eupneic 3-phase respiratory pattern (inspiration, post-inspiration, and stage 2 of expiration). Moreover, KF neurons are required for the adjusting breathing characteristics across a range of conditions.

Much of this evidence derives from studies featuring significant perturbations in KF activity. For example, lesions of the KF substantially alter the respiratory pattern, introducing augmented variability in frequency and amplitude, or even eliminating the post-inspiratory phase and thus disrupting eupneic breathing (Oku and Dick, 1992; Fung and St John, 1995; Dutschmann and Herbert, 2006; Bautista and Dutschmann, 2014a; Bautista et al., 2014; Silva et al., 2016; Jenkin et al., 2017). Chemical excitation of KF neurons enhances constrictor activity in upper airway muscles and prolongs the duration of the post-inspiratory phase of the respiratory cycle (Dutschmann and Herbert, 2006). In contrast, disruption of GABAergic transmission in the KF induces periodic apneas, respiratory irregularities, and the loss of abdominal expiratory contractions (or active expiration) during exposure to high levels of carbon dioxide (Abdala et al., 2016; Dhingra et al., 2016; Barnett et al., 2018). KF neurons are also targets for neuromodulators that modify neuronal excitability and change the breathing pattern and rhythm. In this regard, activating 5-HT_1A_ (serotonin) receptors in the KF reduces spontaneous apneas, while antagonizing these receptors can destabilize breathing frequency (Dhingra et al., 2016). Some KF neurons express *µ*-opioid receptors that, when activated, can cause breathing irregularities and bring about life-threatening apneas, a mechanism associated with opioid-induced respiratory depression (Bachmutsky et al., 2020). In addition to the control of eupneic breathing, the KF also contributes to the respiratory adjustments related to behaviors such as vocalization, swallowing, and coughing (Dutschmann and Dick, 2012; Jakus et al., 2008; Bautista and Dutschmann, 2014b), acting as a convergent and integrative synaptic station that relays inputs from suprapontine regions to the rCPG (Hayward et al., 2004). Despite the evidence showing the critical role of the KF in breathing control, its cellular organization and interaction with other respiratory compartments to adjust the respiratory rhythm and pattern remain uncertain.

There is a long tradition of using mathematical modeling and computational techniques to build theoretical models of the rCPG, develop intuition about how it functions, and test hypotheses on respiratory rhythm and pattern generation (Lindsey et al., 2012; Molkov et al., 2017; Phillips and Rubin, 2022). In the past, we developed models integrating a medullary CPG circuit with a pontine component using an established reduced, activity-based mathematical framework (Barnett et al., 2018; Wittman et al., 2019; Molkov et al., 2017; Flor et al., 2020; Rubin et al., 2009, 2011). Previous models differed in their assumptions about pontine interactions and intrinsic dynamics, which need to be better characterized experimentally. These models have provided a proof of principle for the proposed central role of the KF in respiratory control and have yielded preliminary predictions about interactions of the KF with other pontine and medullary respiratory areas as well as their contribution to respiratory rhythmicity and pattern formation under a variety of conditions. The details of these predictions, however, depend on the assumptions made about intrinsic properties and activity patterns of KF neurons, synaptic interactions in the circuit, and conditions leading to aberrant KF activity and subsequent perturbations of breathing. To reproduce specific experimental observations, model networks that include the KF have required the presence of specific synaptic pathways, such as from inhibitory neurons in the Bötzinger Complex (BötC) (Wittman et al., 2019) and NTS pump cells (Molkov et al., 2013). Models have also suggested that KF neurons may have certain intrinsic properties modulating their activity such as endogenous bursting (Wittman et al., 2019). This capability, however, has never been demonstrated experimentally.

Understanding the mechanisms underlying the control of KF activity is essential to identify its role in breathing pattern formation in eupnea and to determine how changes in these mechanisms affect the expiratory motor pattern and contribute to inducing stereotyped breathing patterns such as those observed in Rett syndrome (Abdala et al., 2016). In this study we consider various KF circuit organizations paired with different baseline activity patterns and local input sources to assess which may explain a set of experimental observations not previously addressed. Specifically, each model configuration was tested against the following list of benchmarks based on the literature: (1) reduction of GABA/5-HT_1A_ inhibition in the KF leads to an aberrant respiratory pattern in which breathing is intermittently disrupted by periods of apnea (Abdala et al., 2016; Dhingra et al., 2016); (2) the duration of apneic periods is relatively constant or increases as the inhibition level to KF decreases (Abdala et al., 2016; Dhingra et al., 2016); (3) when KF is dysfunctional, a sufficient inchibition of the KF neurons transforms intermittent breathing into eupnea (Abdala et al., 2016, 2010); (4) a further increase in inhibition of the KF neurons causes abdominal expiratory activity during the late part of the E2, or late-E, phase of the respiratory cycle, with a frequency that increases in a step-wise, or quantal, manner as inhibition is strengthened (Koolen, 2021). We discuss two viable model configurations, while presenting evidence against certain others, and formulate experimentally testable implications in each case.

## 2 Methods

To study the contribution of KF activity to the generation and modulation of various respiratory patterns, we developed and analyzed two families of computational models of a respiratory brain stem neuronal circuit. In the first family, which we call the *tonic model*, the KF population is homogeneous with steady, sustained or *tonic* activity under normal conditions. The other family combines two KF populations, one that shows sustained tonic activity (as in the tonic model) and another that is intrinsically silent under normal conditions. Based on the inclusion of the second population, we refer to this as the *silent model*. Schematic diagrams of the models are shown in Figure 1.

**Figure 1:**
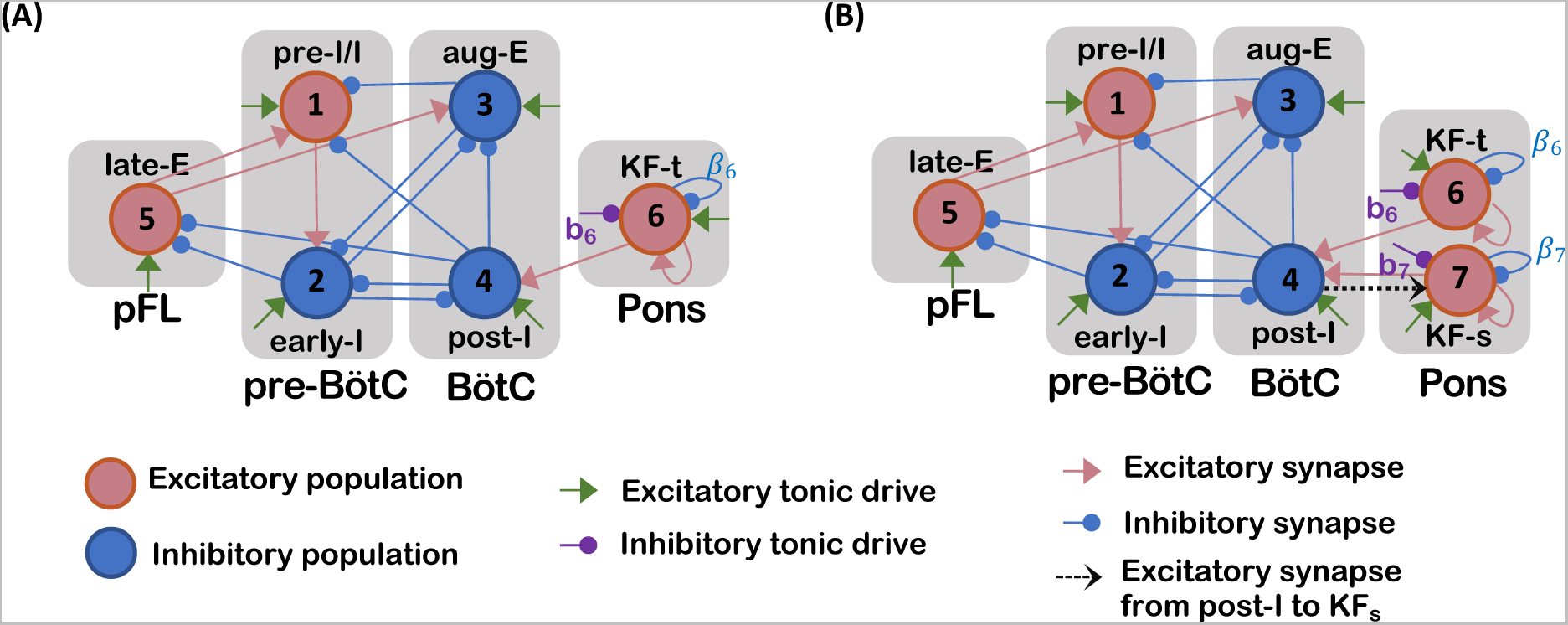
Schematic diagrams for the respiratory network models considered in this work. (A) Tonic model. (B) Silent model. The labels *b*_6_*, b*_7_*, β*_6_*, β*_7_ refer to parameters used to represent the strengths of certain connections explored in the model. Abbreviations referring to neuronal populations and sites are described in the text.

The models presented in this work are adapted from previous models for respiratory neurons and circuits (Rubin et al., 2009, 2011; Wittman et al., 2019). To form these models, we incorporated the KF component into a pre-existing model presented by Rubin et al. (2011). Similarly to that model, the models in this work include the following respiratory populations, which together comprise what we call the *respiratory core*: pre-inspiratory (pre-I) and early-inspiratory (early-I) populations of the pre-Bötzinger Complex (pre-BötC) and post-inspiratory (post-I) and augmenting expiratory (aug-E) populations of the Bötzinger Complex (BötC). The pre-I and early-I populations are active during the inspiratory phase of respiration (I phase), whereas aug-E and post-I are active during the expiratory phase (E phase). More precisely, post-I units feature a surge of activity at the onset of expiration, followed by a gradual tailing off of activity. In contrast, aug-E units gradually ramp up their activity, in an augmenting pattern, over the course of expiration. The models also include a late-expiratory (late-E) population of the lateral parafacial region (pFL), which remains inactive during resting breathing (Bianchi et al., 1995).

All of the synaptic connections between these respiratory populations in the models shown in Figure 1 are either directly shown in, or inferred from, experimental observations (Rubin et al., 2011). Both models also include a KF sub-population, named KF-t, that is tonically active under normal conditions and helps to maintain eupnea. The silent model includes KF-t and another KF sub-population, named KF-s, that is silent under normal conditions. The interactions of the KF units with the respiratory core in these models occur through excitatory connections from KF-t and KF-s to the post-I population. These connections have been proposed previously (Molkov et al., 2013; Jenkin et al., 2017; Barnett et al., 2018; Geerling et al., 2017). Little is known about sources of inhibition to KF neurons. For exploratory purposes, we considered both tonic inhibition to the KF originating from an unspecified outside source and recurrent inhibitory connections within the KF. We analyze the extent to which each of these inhibition types, within each model, yields outputs consistent with respiratory perturbations seen in Rett syndome (RTT), and we also study the emergence of late-E activity (active expiration). Previous work by Wittman et al. (2019) introduced a simplified representation of pulmonary stretch receptor feedback related to I phase output. As part of the analysis of the silent model done in the current paper, we study the primary effect of this pathway by considering what happens if we introduce an additional excitatory connection from post-I to KF-s (shown as a black dashed arrow in Figure 1B). Although this connection would need to run through another inhibitory population and thus be manifested via disynaptic disinhibition in reality, for simplicity we treat it as a monosynaptic excitation in this work.

Each component in these models is described by a single two-dimensional system of Hodgkin-Huxley type equations representing the dynamics of a membrane voltage variable *v* and a secondary slow variable *h* or *m*. We consider this framework as a simplified representation of dynamics in which the activity levels of neurons within each population are synchronized, and the reduction that it provides allows us to use the tools of phase plane analysis to understand model dynamics and the effects of parameter variations. To present the equations for this system, we assign a subscript from 1 to 5 for the pre-I/I, early-I, aug-E, post-I and late-E units, respectively. The Kf-t unit is labeled as 6 and the additional Kf-s unit in the silent models as 7.

The pre-I/I (*i* = 1) and late-E (*i* = 5) units feature a persistent sodium current (Butera et al., 1999; Rubin et al., 2011) and hence are represented by the following differential equations:

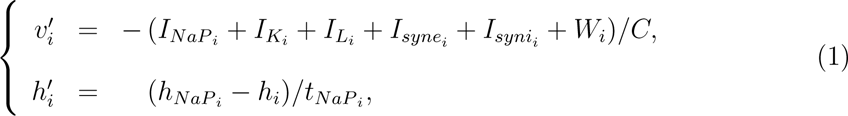

where *I_NaPi_* is the persistent sodium current, *I_Ki_* represents the delayed rectifier potassium current, *I_Li_* is the leak current, *I_synei_* and *I_synii_* are the excitatory and inhibitory synaptic currents, and *W_i_* represents the noise, respectively, for each unit. All of the other units in the model network (*i* ∈ {2, 3, 4, 6, 7}) are modeled as adaptive neurons and obey the following equations:

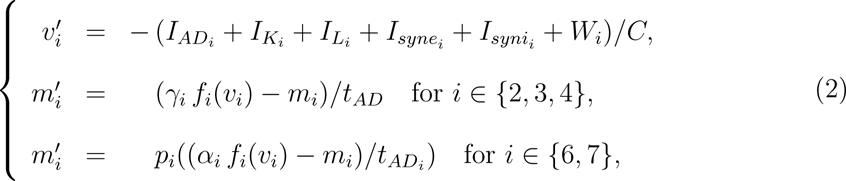

where *I_ADi_* is a second potassium current causing adaptation during spiking and *t_AD_* is a constant shared for all *i* ∈ {2, 3, 4}. Importantly, the parameters *t_NaP i_* from (1) and 1*/t_AD_* and *p_i_/t_ADi_* from (2) are small relative to the timescale of the voltage variables, such that the *h_i_* and *m_i_* evolve relatively slowly.

The various currents in (1) and (2) are defined as follows:

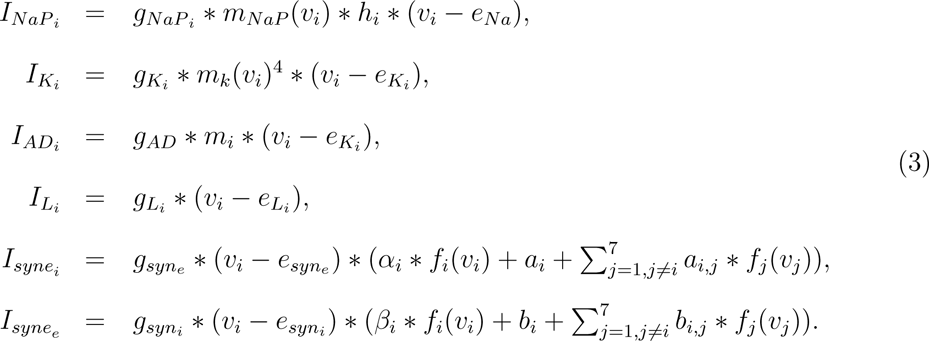

In system (3), we introduce asterisks to indicate multiplication, to avoid confusion relating to the *v_i_*-dependent terms in the first two equations.

The noise term, *W_i_*, in (1) and (2) is

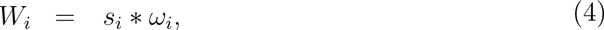

where *ω_i_* ∼ N (0, 1) (i.e., a normally distributed random variable with mean 0 and standard deviation 1). For most of the analysis in this work, we turn off the noise by setting *s_i_* = 0. We indicate clearly where noise has been turned on when we describe the results.

The activation functions associated with (1),(2), and (3) are given by:

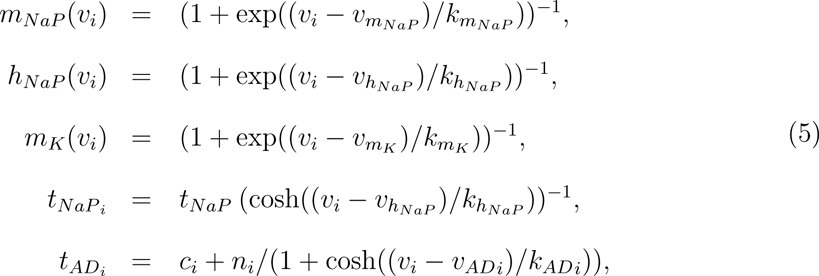

with the constants *c_i_, n_i_* defined for *i* ∈ {6, 7}. The *output function f_i_* for *i* ∈ {1, 2, 3, 4, 5}, which appears in both (2) and (3), takes the piecewise linear form

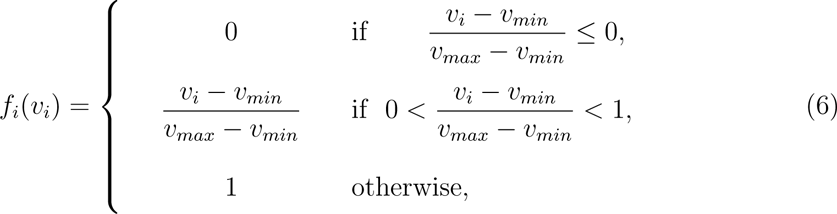

whereas the output function *f_i_* for *i ∈ {*6, 7*}* is given by

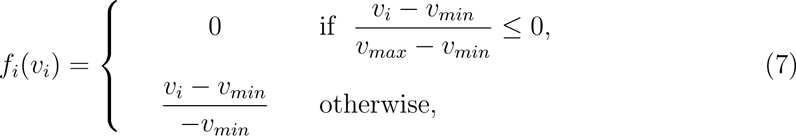

for constants *v_min_* and *v_max_*. The default values of these constants and all other parameters used in the models appear in Tables 1, 2.

**Table 1:** Default parameter values for the tonic model :

|  |  |  |  |  |  |  |  |
| --- | --- | --- | --- | --- | --- | --- | --- |
| $g_{Nap_i}$ | 5.0 nS ($i \neq 5$) | $g_{Nap_5}$ | 4.72 nS | $g_{K_i}$ | 5.0 nS ($i \neq 6$) | $g_{K_6}$ | 0.0 nS |
| $g_{L_i}$ | 2.8 nS ($i \neq 6$) | $g_{L_6}$ | 2.5 nS | $g_{AD}$ | 10.0 nS | $g_{syn_e}$ | 10.0 nS |
| $g_{syn_i}$ | 60.0 nS | $e_{Na}$ | 50.0 mV | $e_{K_i}$ | -85.0 mV ($i \neq 6$) | $e_{K_6}$ | -90.0 mV |
| $e_{L_i}$ | -60.0 mV ($i \neq 6$) | $e_{L_6}$ | -66.5 mV | $e_{syn_e}$ | 0.0 mV | $e_{syn_i}$ | -75.0 mV |
| $a_1$ | 0.03 | $a_2$ | 0.875 | $a_3$ | 0.9 | $a_4$ | 0.6 |
| $a_5$ | 0.11 | $a_6$ | 0.15 | $a_{12}$ | 0.5 | $a_{51}$ | 0.5 |
| $a_{53}$ | 0.25 | $a_{64}$ | 0.95 | $\alpha_6$ | 1.0 | $b_6$ | 0.001 |
| $b_{23}$ | 0.42 | $b_{24}$ | 0.22 | $b_{25}$ | 0.09 | $b_{31}$ | 0.15 |
| $b_{32}$ | 0.1 | $b_{41}$ | 1.0 | $b_{42}$ | 0.66 | $b_{43}$ | 0.2 |
| $b_{45}$ | 0.101 | $\beta_6$ | 0.05 | $t_{NaP}$ | $4.0 \times 10^3$ | $t_{AD}$ | $2.0 \times 10^3$ |
| $vm_{NaP}$ | -40.0 mV | $vm_K$ | -30.0 mV | $vh_{NaP}$ | -55.0 mV | $v_{max}$ | -20.0 mV |
| $km_{NaP}$ | -6.0 mV | $km_K$ | -4.0 mV | $kh_{NaP}$ | 10.0 mV | $v_{min}$ | -50.0 mV |
| $c_6$ | $7.0 \times 10^2$ | $n_6$ | $1.0 \times 10^4$ | $v_{AD6}$ | -42.0 mV | $k_{AD6}$ | 0.9 mV |
| $\gamma_i$ | 1.0 ($i \neq 4$) | $\gamma_4$ | 2.0 | $s_i$ | 0.0 ($\forall i$) | $p_6$ | 0.0286 |

**Table 2:** Default parameter values for the silent model. The other parameter values for the model are as given in Table 1

|  |  |  |  |  |  |  |  |
| --- | --- | --- | --- | --- | --- | --- | --- |
| $a_7$ | 0.1 | $a_{74}$ | 0.75 | $\alpha_7$ | 1.0 | $\beta_7$ | 0.0 |
| $n_7$ | $5.0 \times 10^3$ | $v_{AD7}$ | -50.0 mV | $k_{AD7}$ | -0.5 mV | $p_7$ | 0.02 |
| $b_7$ | 0.02 | $c_7$ | $4.0 \times 10^2$ | | | | |

## 3 Results

### 3.1 Tonic Model

#### 3.1.1 Transitions from normal breathing to periodic breathing

For the default parameter values given in Table 1, the tonic model generates a eupneic breathing pattern (Figure 2A). Notice that KF-t is assumed to exhibit tonic activity, with a constant non-zero output *f*_6_(*v*_6_) ≈ 0.15 (pink), which may result from sustained drive from elsewhere in the pons that may potentially be tuned by feedback pathways. The tonic excitatory input from KF-t promotes the long E phase duration compared to the I phase duration during normal breathing activity in the model. Experiments show that a significant reduction of the GABA inhibition to the KF leads to respiratory apneas (Abdala et al., 2016), suggesting the hypothesis that such a reduction underlies respiratory disruptions as seen in RTT. When the level of sustained inhibition to KF-t in the model is reduced, KF-t produces endogenous oscillations, which in turn lead to a prolonged active phase of the post-I population on some cycles that interrupt the normal breathing pattern and would manifest as respiratory apneas in a physiological system (Figure 2B). A comparison of the KF-t output traces between the two panels of Figure 2 shows that while KF-t output is elevated during the active phase of its oscillations relative to its sustained output level in the tonic case, the KF-t output becomes lower than the tonic level during the silent phases, or inter-burst intervals, in the oscillatory case. This difference causes the respiratory cycles that occur in between apneas within periodic breathing (Figure 2B) to be shorter than the normal cycle periods seen in eupneic breathing (Figure 2A), consistent with experimental observations under RTT-like conditions (Abdala et al., 2016).

**Figure 2:**
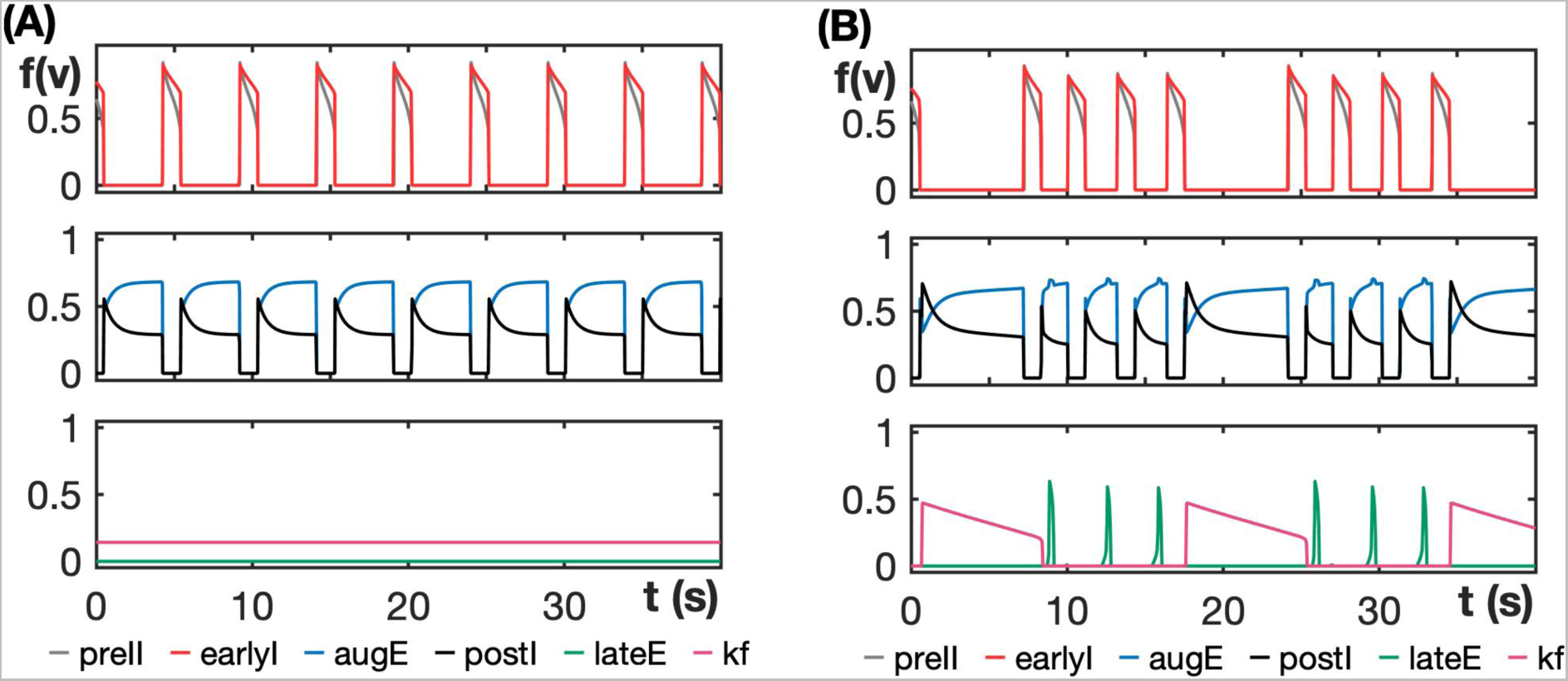
Tonic model output patterns. (A) Normal respiratory rhythm for the default parameter values given in Table 1. (B) Periodic breathing when recurrent inhibition is shut off, *β*_6_ = 0.0 (simulated RTT). KF engages in slow oscillations (pink trace, bottom subplot) that drive prolonged E phases.

We used nullcline analysis to better understand which forms of inhibition to KF-t could support the generation of the activity patterns that we expect based on experimental findings and to explain why the tonic KF-t output switches to an oscillatory pattern when inhibition is reduced in a sustained way. We see in Figure 3 that KF-t, as modeled by (2), has a cubic *v*_6_-nullcline (solid blue). The *m*_6_-nullcline (solid red) intersects the active branch of the *v*_6_-nullcline for default parameter values, corresponding to a stable equilibrium point at *v*_6_ ≈ −42 (solid grey square). Notice in Figure 3A that when we reduce the recurrent or self-inhibition within the KF-t (lower *β*_6_), the slow nullcline intersects the middle branch of the *v*_6_-nullcline, destabilizing the equilibrium point and leading to oscillatory behavior of KF-t. The oscillatory trajectory of the KF-t unit in this regime is overlaid in grey in the figure panel. The trajectory oscillates between the left and right stable branches of the fast *v*_6_-nullcline. The reduction of tonic inhibition (*b*_6_) that comes in from outside to the KF-t, in contrast, does not depend on the output of the KF-t unit and hence shifts the *v*_6_-nullcline vertically, which does not induce a transition in the stability of the equilibrium point and hence does not lead to KF-t oscillations (Figure 3B). Thus, we henceforth assume for this model that the recurrent inhibition strength *β*_6_ is non-zero in the eupneic regime and decreases in the RTT condition, which represents a prediction of this work, and for simplicity we assume that the tonic inhibition strength *b*_6_ = 0, since this form of inhibition is not necessary to explain experimental observations.

**Figure 3:**
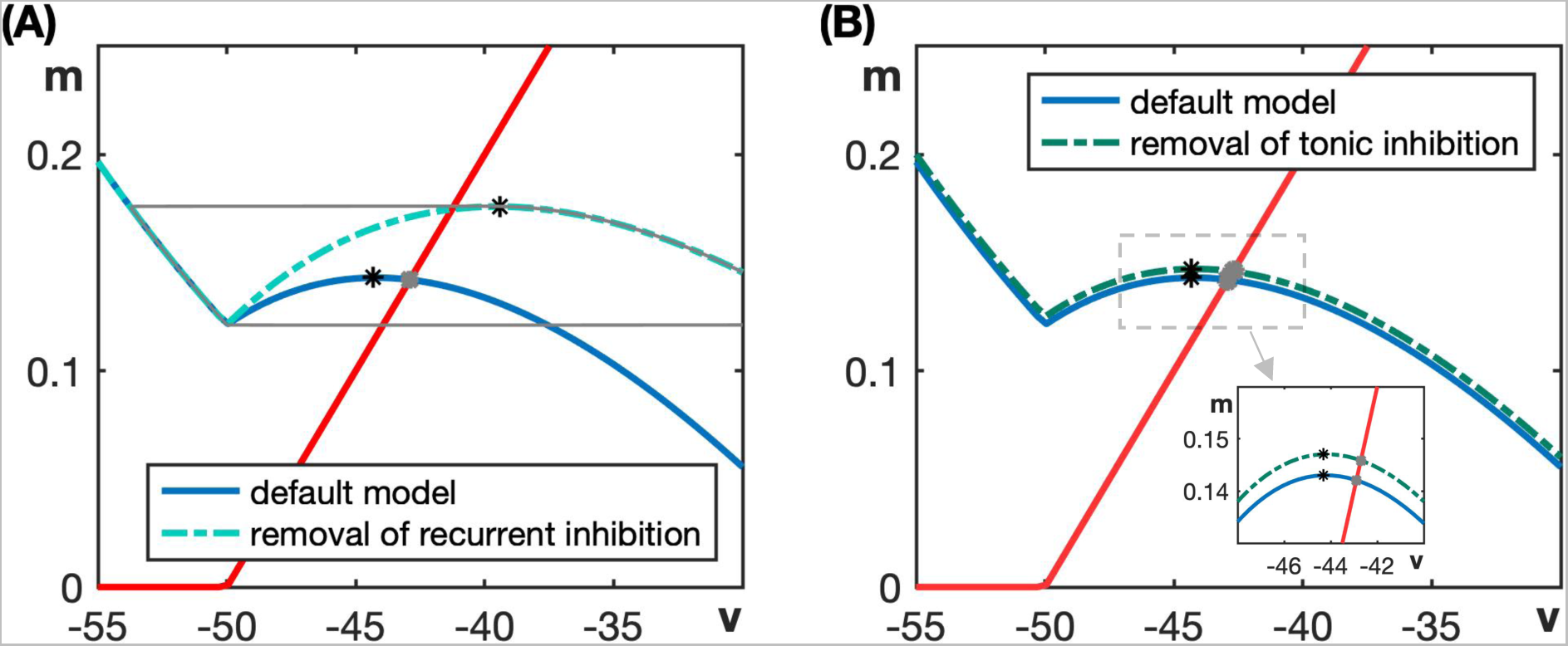
Tonic KF-t model in the phase plane. (A) The default model has a stable equilibrium point with *v*_6_ ≈ −42 (grey square), corresponding to tonic activity. When the recurrent inhibition to KF-t is removed, the local maximum (black asterisk) of the resulting *v*_6_ nullcline (blue dashed curve) shifts to the right of the slow nullcline (red curve). Oscillations emerge in KF-t in this case (grey orbit). (B) The *v*_6_ nullcline shifts upwards when tonic inhibition is removed as shown by the dashed blue curve. KF-t remains tonic in this case since the local maximum of the *v*_6_ nullcline (black asterisk) remains on the left of the equilibrium point (see also zoomed view in inset).

While Figures 2, 3 illustrate the extreme case of full removal of the inhibition to KF-t in the tonic model (*β*_6_ = 0), Figure 4A,B show the periodic breathing exhibited by the tonic model at the partially reduced inhibition levels *β*_6_ = 0.025, which is near the value where KF-t transitions from tonic to oscillatory behavior, and *β*_6_ = 0.015. For these intermediate inhibition levels, although the KF-t equilibrium point lies on the middle branch of the *v*_6_-nullcline and is unstable, its position is in between the two intersection points shown in Figure 3A; that is, it lies closer to the local maximum of the *v*_6_-nullcline. The proximity of the equilibrium and the *m*_6_-nullcline to this maximum means that the trajectory evolves very slowly at the end of the KF-t active phase, resulting in a longer KF-t oscillation period and larger duty cycle for larger *β*_6_. This effect explains the prolonged KF-t oscillations in Figure 4A, B.

**Figure 4:**
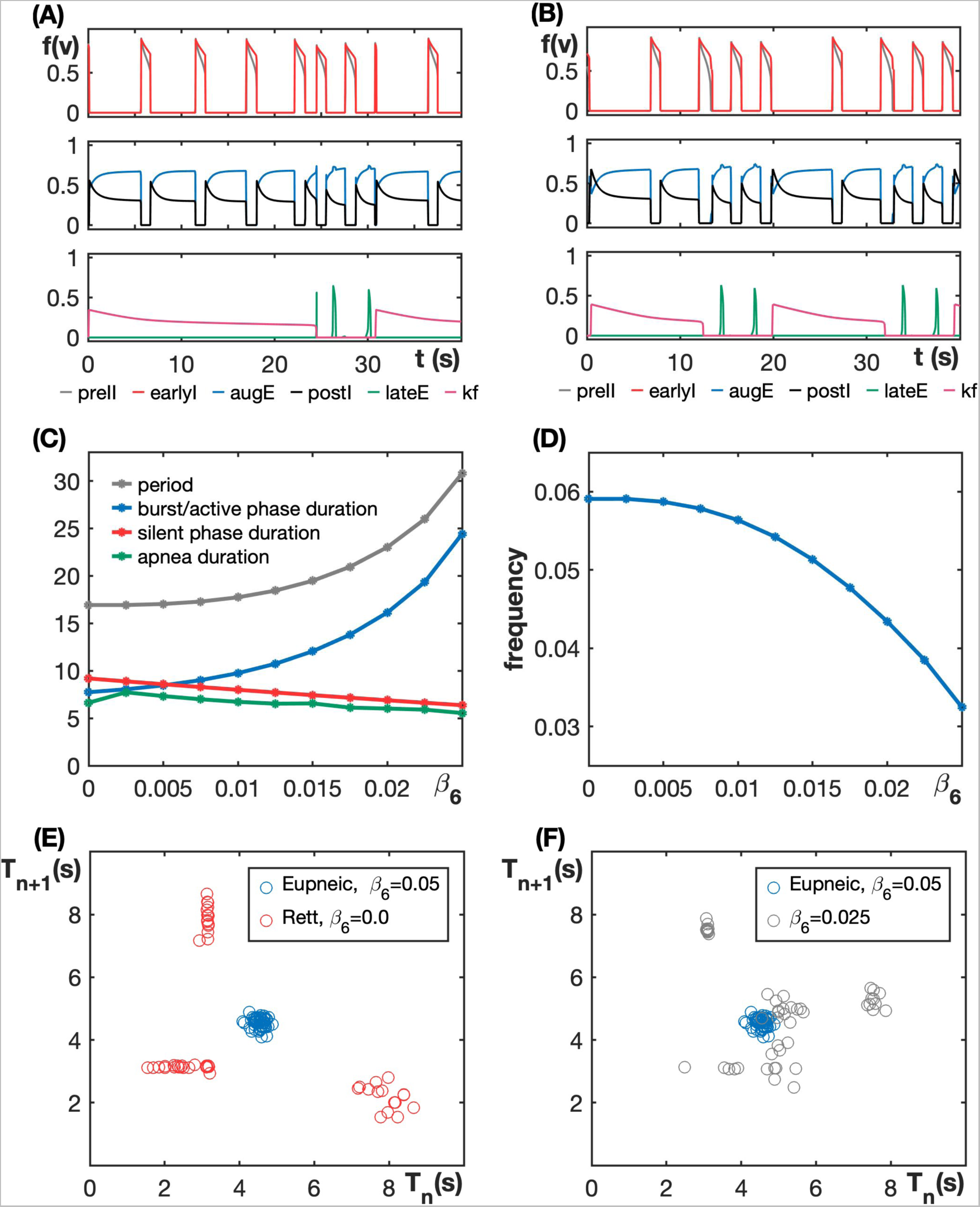
Tonic model output depends on the level of recurrent inhibition within the KF-t population (*β*_6_). (A) Respiratory pattern with a recurrent inhibition strength near the point of transition of KF-t from tonic to oscillatory (*β*_6_ = 0.025). (B) Respiratory pattern with recurrent inhibition strength *β*_6_ = 0.015, representing an intermediate case between that shown in (A) and the RTT condition shown in Fig. 2B. (C) When *β*_6_ is decreased, the period of KF-t oscillations and its burst or active phase duration decreases while apnea duration remains roughly the same but slightly increases. (D) Apnea frequency, defined as KF-t burst frequency (i.e., the inverse of the period of oscillations), rises as the inhibition level to KF-t decreases. (E) Total respiratory cycle duration during the (*n*+1)*^st^* cycle relative to the *n^th^* cycle for the tonic model with noise, *β*_6_ = 0.05 (default value, blue dots) and *β*_6_ = 0.0 (RTT, red dots). (F) Total respiratory cycle duration during the (*n* + 1)*^st^* cycle relative to the *n^th^* cycle for the tonic model with noise, *β*_6_ = 0.05 (default value, blue dots) and *β*_6_ = 0.025 (gray dots).

Even though the KF oscillations are longer for these transitional values of *β*_6_ than for the simulated RTT condition (*β*_6_ = 0), notice that the actual apnea durations are relatively short for these values and increase slightly as *β*_6_ decreases (Figure 4C). How can longer KF-t active durations produce shorter apneas when apneas in the model result from KF-t drive to the post-I unit? The key to explaining this outcome is to focus on the level of KF-t output while it is active. Our simulations show that when KF-t output is above about 0.23, it supports prolonged post-I activity corresponding to apneas. When KF-t output falls below this value, closer to levels comparable to the output in the tonic regime with *β*_6_ = 0.05, the normal respiratory rhythm takes over. For intermediate *β*_6_ near the transition to KF-t oscillations, during each prolonged KF-t active period, KF-t output relatively quickly reaches close to the tonic level and plateaus. Thus, apnea duration is relatively short. When *β*_6_ is reduced from there, although KF-t active phases become shorter, the amplitude of each oscillation increases (see Figure 3A) and it takes longer for KF-t output to decrement to tonic levels. Therefore, even though the KF-t burst duration decreases when inhibition is lowered, the apnea duration increases. As a final subtlety, we note that the actual apnea duration plotted in Figure 4C (green curve) is not monotonic. This non-monotonicty arises because the apnea duration also depends on the phase of post-I activity when KF-t becomes active. Regardless of this phase, post-I stays active for a similar amount of time after KF-t activation occurs, based on the level of KF-t output. Thus, when KF-t activation occurs relatively late within the E phase, the overall apnea will be long, and this phase relation need not vary monotonically with *β*_6_.

Since KF-t output falls to near tonic levels during each KF-t active phase, we only count the first post-I active period within each of these KF-t cycles as an apnea. Thus, the frequency of apneas increases as the recurrent inhibition *β*_6_ is lowered from 0.025 (Figure 4D). These findings match the experimental results reported previously (Abdala et al., 2016; Koolen, 2021).

To further study burst patterning and transitions in activity patterns when recurrent inhibition to KF-t is lowered, we modified the tonic model to include additive Gaussian noise in the voltage equation (*s_i_* = 1.0 in (1) and (2)), as described in Section 2. Figure 4E,F show the total respiratory duration (i.e., from one E onset to the next) during the (*n* + 1)*^st^* burst relative to the duration for the *n^th^* burst (*T_n_*_+1_ versus *T_n_*) for inhibition levels *β*_6_ = 0.0 (RTT) and *β*_6_ = 0.025 (just after the onset of KF-t oscillations) along with the same information for the default value *β*_6_ = 0.05. During normal breathing (blue dots), this level of noise induces little variability in the respiratory period (which takes values between 4 and 5*s*). In the case of RTT (red dots), respiratory periods cluster around two values, one corresponding to long apneas each having a duration of nearly 8*s* and the other to shorter normal breathing periods of approximately 3*s*. The reason that the normal, non-apneic respiratory bursts in RTT are shorter than those defined for the default model parameters is that, as we pointed out with respect to Figure 4, the inter-burst KF output level within KF-t oscillations is much lower than its tonic output level at the baseline level of inhibition.

For the sub-normal inhibition level *β*_6_ = 0.025 (grey dots) considered in Figure 4F, the durations of the longest post-I cycles are generally shorter than the prolonged apneas in the RTT case (Figure 4F, red dots), as discussed above. In the intermediate case, as we have also discussed already, each post-I cycle duration depends on the gradually declining KF-t output level (Figure 4A), and hence there is much more variability in the respiratory cycle duration, with less of a clear clustering of durations, than for the extreme *β*_6_ values. These results are consistent with the findings in (Abdala et al., 2016). In particular, Figure 5Cb in (Abdala et al., 2016) shows that the effects of blocking KF GABA_A_ receptors in wild-type rats are comparable to Figure 4E.

Past literature has shown that a systemic application of various 5-HT_1A_R agonists can reduce the frequency of spontaneous apneas and restore normal respiratory function to various degrees in the murine model of RTT (Abdala et al., 2010; Dhingra et al., 2016). These agonists can potentially boost a potassium current and thus enhance tonic inhibition within the KF (Levitt et al., 2013). They can also facilitate glycinergic neurotransmission (Shevtsova et al., 2011) or opening of chloride channels (Manzke et al., 2010), each of which may underlie an increase in recurrent inhibition. In Figure 5A, we show nullclines for the KF-t unit, comparing the RTT scenario in which KF-t is oscillatory to a case with increased recurrent inhibition. The slow nullcline (solid red) intersects the middle branch of the *v*_6_-nullcline. This comparison repeats that given in Fig. 3A, showing that if recurrent inhibition to KF-t (*β*_6_) increases, the tonic KF activity is restored, which implies that periodic breathing will switch back to the normal breathing pattern. On the other hand, from the RTT condition, increasing the external tonic inhibition (*b*_6_) to KF-t does not switch the nullcline intersection off of the middle branch (Fig. 5A, dashed *v*_6_-nullcline) and hence maintains the oscillatory KF activity, until with a sufficient increase in *b*_6_ the intersection moves to the left branch of the *v*_6_-nullcline (Fig. 5A, dash-dotted *v*_6_-nullcline) and stabilizes. This stable equilibrium point location implies that KF-t becomes fully inactive and no longer supports normal breathing.

**Figure 5:**
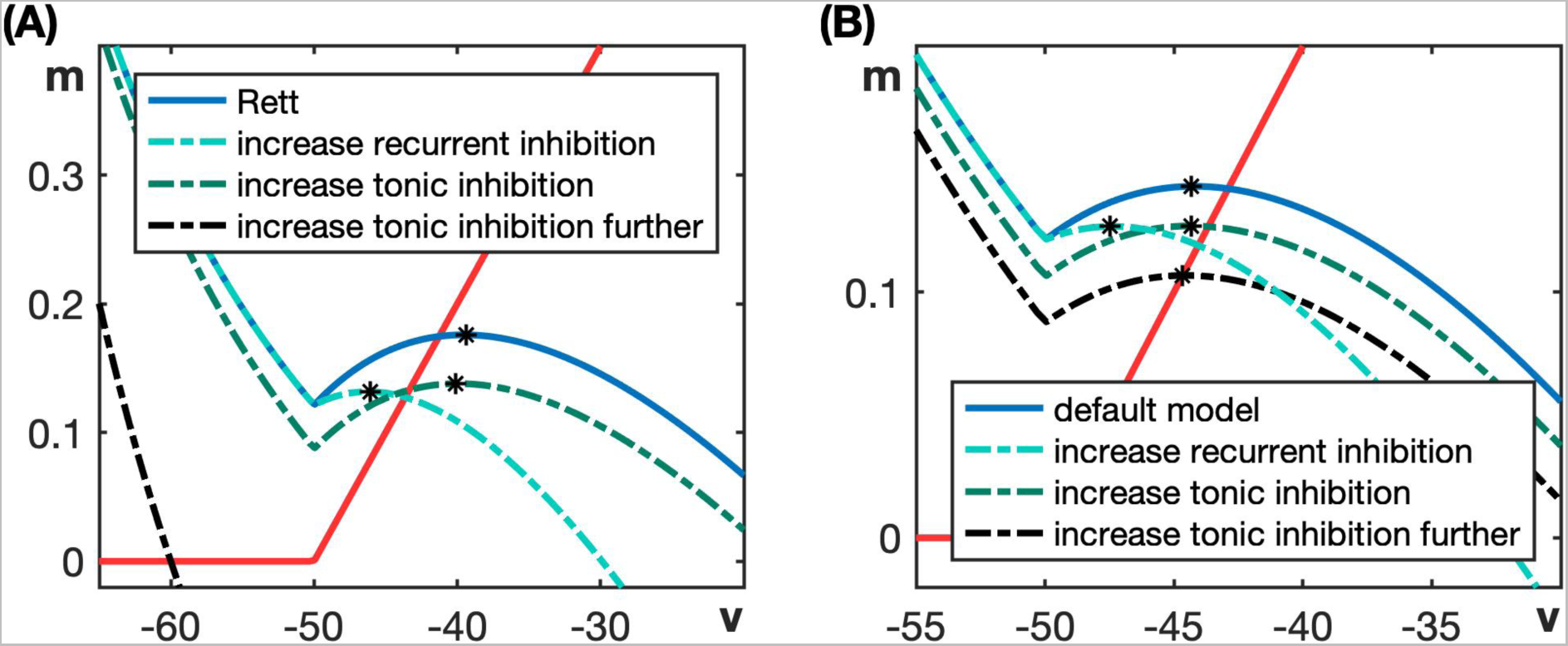
(A) Starting with oscillatory KF-t (RTT), *β*_6_ = 0.0. When recurrent inhibition, *β*_6_, is increased from 0.0, KF-t becomes tonic. When tonic inhibition, *b*_6_, is increased, KF-t remains oscillatory. When it’s increased further, it becomes silent. (B) Tonic model with inhibition increased above baseline. When recurrent inhibition is increased, KF-t remains tonic with its equilibrium point on the right branch of the *v*_6_-nullcline but the point moves to a lower *v*_6_ value, corresponding to less KF output. When tonic inhibition is increased, the equilibrium point moves to the middle branch and destabilizes, and thus a transition from tonic KF output to KF oscillations occurs.

Overall, these simulations indicate that a reduction in the recurrent inhibition to the KF neurons that provide tonic input to the ventromedullary neurons can trigger episodic apneas and drive breathing irregularities. In this scenario, reduction of tonic inhibition to the KF does not produce similar effects. A prediction emerging from this analysis is that drugs that boost endogeneous inhibition would be more efficacious at reversing breathing irregularity than drugs that boost or produce an exogenous tonic inhibitory drive.

#### 3.1.2 Emergence of active expiration

Expiration is a process that becomes active in conditions of increased respiratory drive. Exposure to low oxygen (hypoxia) or high carbon dioxide levels (hypercapnia) triggers active expiration with the recruitment of abdominal muscles during the late part of the E2 phase (i.e., late expiration, or late-E) to improve pulmonary ventilation (Baertsch and Ramirez, 2019). Decreased KF activity is known to play a role in the emergence of late-E abdominal activity (Jenkin et al., 2017; Barnett et al., 2018). Moreover, Koolen (2021) showed that systemic administration of the 5-HT_1A_R agonist NLX-101 increases the respiratory drive, elevating the respiratory frequency and causing the appearance of active expiration (late-E activity) under resting conditions.

We first consider the phenomenon of late-E activation. The tonic model has an excitatory synaptic connection from KF-t to post-I and an inhibitory connection from post-I to late-E (see Figure 1A). This architecture suggests that when there is a reduction in KF activity, a decrease in post-I output will result, which in turn will reduce the post-I dependent inhibition to late-E. If KF-t output is reduced sufficently, then late-E should start spiking. This logic leads us to consider the input-related mechanisms by which 5-HT_1A_ agonists may strengthen inhibition in the KF. In Figure 5B, we analyze the effect of increasing each of the two different types of inhibition to KF-t. Initially, the KF-t unit has a stable equilibrium at *v*_6_ ≈ −42. When we increase recurrent inhibition, the slow nullcline (shown in red) still intersects the modified *v*_6_-nullcline (shown in dashed blue) on its right branch, leading to a stable tonic equilibrium (Figure 5B). In this case, however, the stable equilibrium has a lower *v*_6_ value than originally. Therefore, we see that increasing recurrent inhibition, *β*_6_, reduces the tonic output level *f*_6_(*v*_6_) of KF-t and eventually causes KF-t output to turn off. If we instead turn on and increase tonic inhibition to KF-t (*b*_6_ *>* 0), notice in Figure 5B that at first a small reduction in the *v*_6_-coordinate of the equilibrium point and hence in *f*_6_(*v*_6_) occurs, but this is not sufficient to promote late-E activity. With an additional increase in *b*_6_, the equilibrium point moves to the middle branch of the *v*_6_-nullcline and KF-t becomes oscillatory, leading to continued late-E suppression. Therefore, to see the emergence of late-E spiking in the tonic model, we gradually increased the recurrent inhibition (*β*_6_) to KF-t and kept *b*_6_ = 0.

For the parameter values given in Table 1, there is no late-E activation, as seen in Figure 2A. An emergence and quantal acceleration of late-E activity with increasing *β*_6_ are illustrated in Figure 6. When we increase *β*_6_ to 0.3 (Figure 6A), late-E spikes once during every three post-I bursts. For *β*_6_ = 0.6 (Figure 6B), late-E spikes once during every two post-I bursts. Increasing *β*_6_ to 1.8 (Figure 6C) results in a late-E spike during every post-I burst. The increasing late-E spiking frequency with respect to *β*_6_ is summarized in Figure 6D. This figure plots the ratio of the number of late-E spikes to the number of post-I bursts on the y axis. For example, at *β*_6_ = 1.2, the ratio is 2*/*3, corresponding to two late-E spikes during every three post-I bursts.

**Figure 6:**
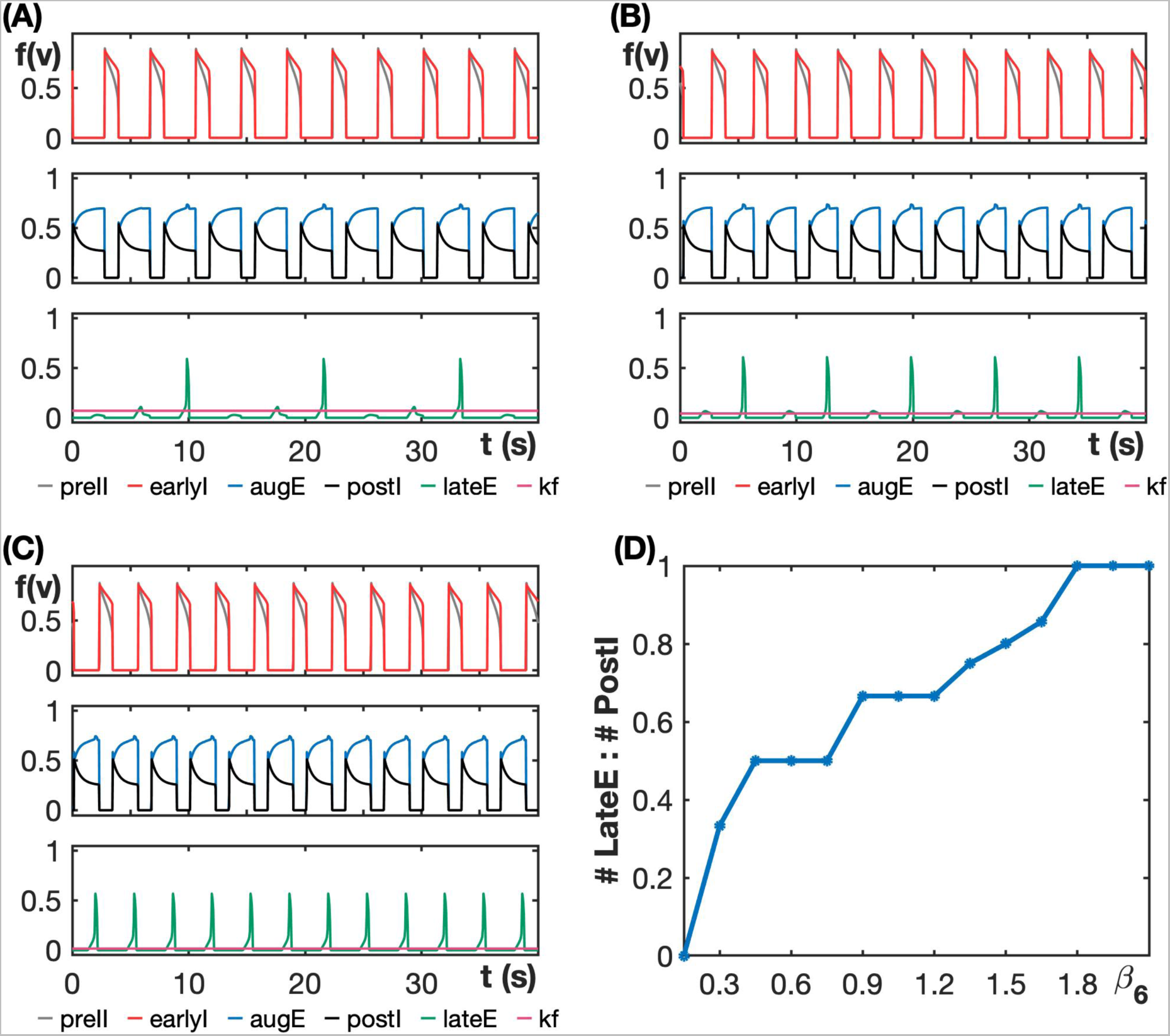
Quantal acceleration of late-E in the tonic model. (A)-(C) Burst patterns exhibited by the tonic model at different inhibition levels. (A) *β*_6_ = 0.3. (B) *β*_6_ = 0.6. (C) *β*_6_ = 1.8. (D) The late-E spiking frequency increases with respect to the recurrent inhibition strength *β*_6_.

In the experiments of Koolen (2021), this increase was largely due to a decrease in the post-I duration while the duration of pre-I oscillations remained roughly the same. Under increases in *β*_6_, our tonic model reproduces these findings; indeed, we can see in Fig. 6 that the E phase becomes shorter as *β*_6_ increases from panel A to panel C (see middle subplots), due to the reduced KF excitation to post-I, while the I phase duration essentially does not change (see top subplots). These results are quantified across a range of *β*_6_ values in Figure 7. Overall, the fact that our model results parallel the data from previous experiments indicates that the assumptions we have made in constructing the tonic model, including the importance of recurrent inhibition within the KF-t populations, are plausible and suggests that our findings may be useful to guide future experiments.

**Figure 7:**
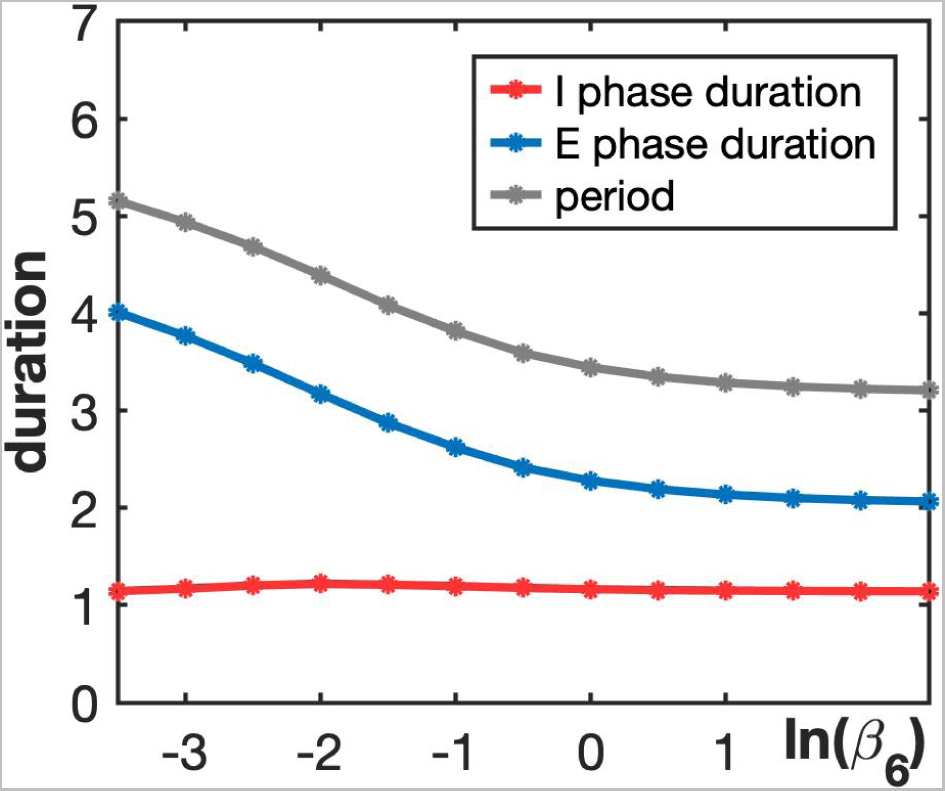
Durations of inspiration and expiration and overall respiratory period vary with ln(*β*_6_) in the tonic model. The gradual decrease in the period with increasing *β*_6_ comes from a drop in the E phase duration, whereas the I phase duration remains roughly constant.

### 3.2 Silent Model

In contrast to the tonic model, the silent model has two KF units: KF-t and KF-s. The values for all of the parameters that appear in both the tonic model and the silent model remain the same as in the tonic model (Table 1), while the values of parameters specific to the silent model are given in Table 2. With these values, KF-t is tonic, with a stable critical point for which *f*_6_(*v*_6_) ≈ 0.15, and KF-s is silent, with a stable critical point below *v_min_* such that its output is zero. Note that the lack of output of the KF-s unit does not reflect an inability to activate; rather, this unit remains quiescent in the baseline model tuning relevant for eupneic rhythm generation, just as the late-E population does. The bursting pattern of the silent model for the default parameter values naturally matches that of the tonic model since KF-s is silent (Figure 8). Hence, our analysis of the role of KF in active expiration resulting from increases in inhibition within the KF, and the corresponding Figures 5B, 6 and 7, apply for the silent model as well.

**Figure 8:**
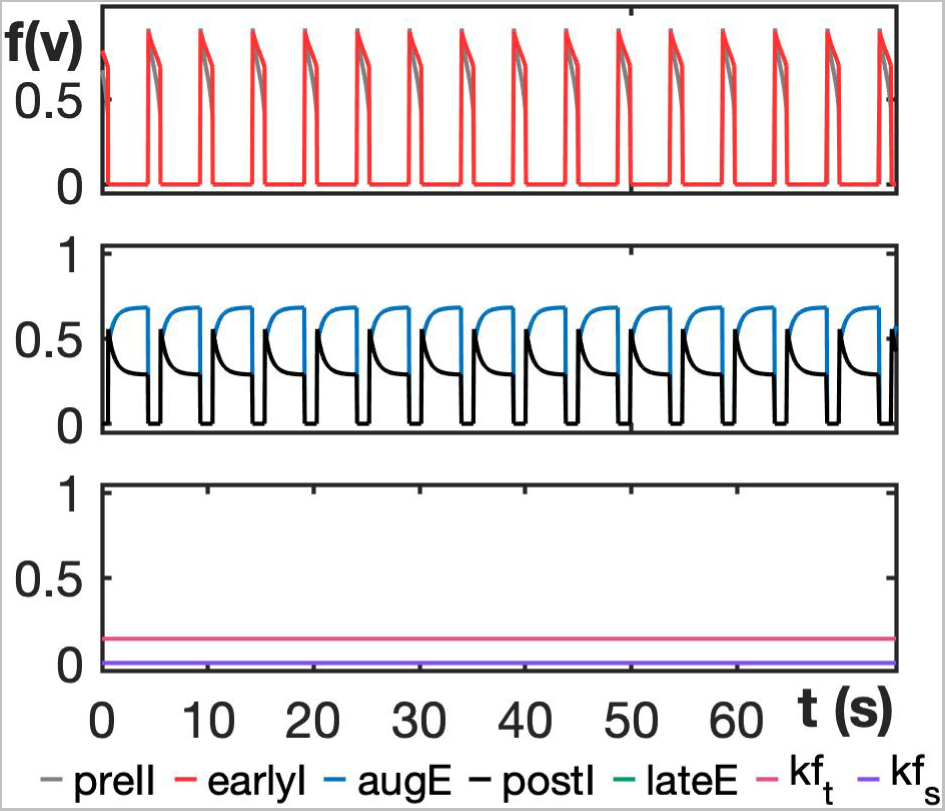
Eupneic respiratory rhythm exhibited by the silent model for default parameter values given in Tables 1, 2.

As with the tonic model, we consider nullclines for the silent model as a means to determine what forms of inhibition of the KF allow the model to match experimental findings on RTT-like respiratory patterns. Importantly, in the silent model, we assume that if the KF-s population becomes active, then it will provide drive to post-I, and in our analysis, we manipulate the inhibition to KF-s. This approach makes the silent model a distinct alternative to the tonic model. If we instead varied inhibition to the KF-t unit in the context of the silent model, then the KF-s unit would simply remain silent and hence would be irrelevant, so our results would trivially match the previous subsection.

Nullcline analysis shows that for the silent model, KF-s transitions to oscillatory behavior when we reduce the tonic inhibition strength to the KF-s unit, *b*_7_ (Figure 9A), because this loss of inhibition causes the model to no longer have a stable fixed point in the silent phase. The same does not happen with recurrent inhibition, since KF-s is initially silent and hence receives no recurrent inhibitory input regardless of *β*_7_; for simplicity, we fix *β*_7_ = 0 in our default parameter set. In the subsequent subsections, we will consider the effects of reduced tonic inhibition in the silent model.

**Figure 9:**
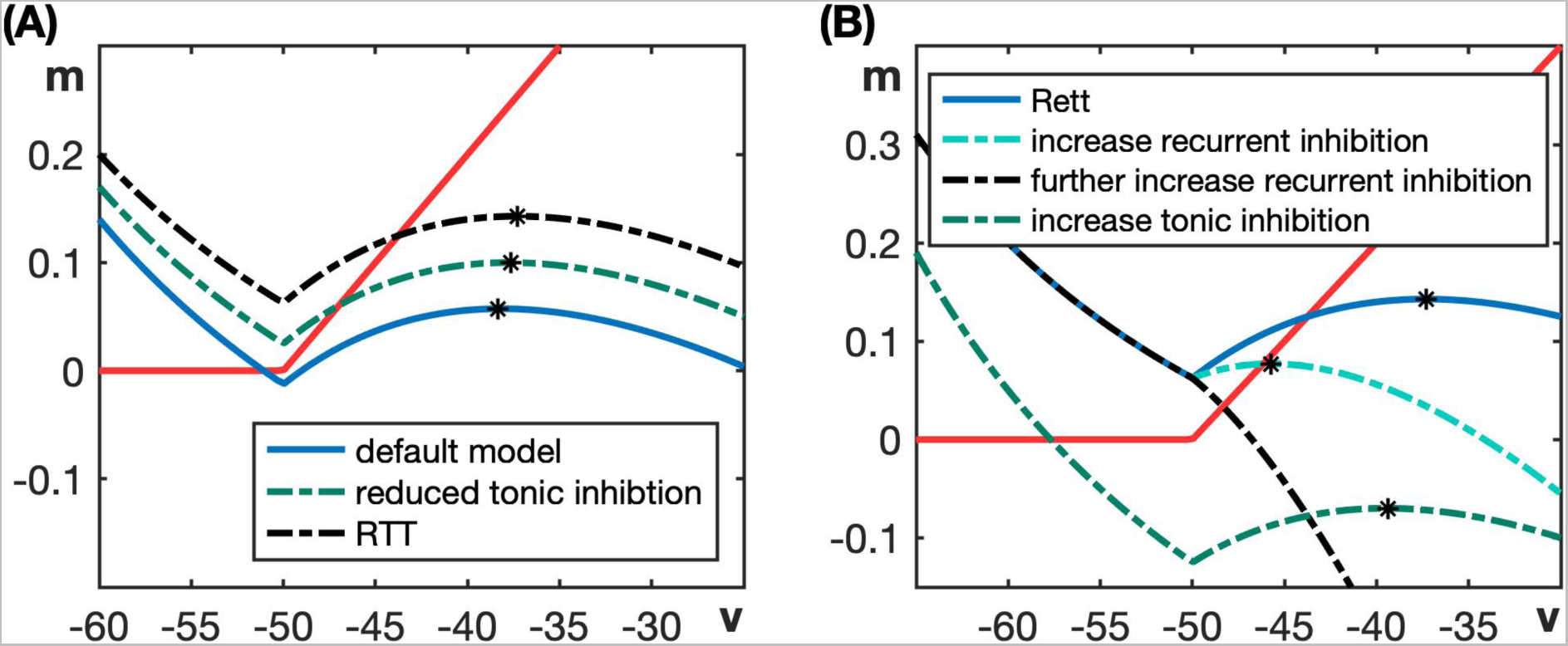
Phase plane analysis of impacts of altered inhibition in the silent model. (A) When the tonic inhibition to KF-s is removed, the slow nullcline (red curve) intersects the middle branch of the resulting *v*_7_-nullcline (green dashed curve). Therefore, oscillations emerge in KF-s in this case. (B) Starting with oscillatory KF-s (RTT scenario), when tonic inhibition is increased, KF-s becomes silent (i.e., nullclines intersect on left branch of dashed green *v*_7_-nullcine). When recurrent inhibition is increased, KF-s remains oscillatory (intersection on middle branch of cyan dashed *v*_7_-nullcline). When it is increased further, KF-s becomes tonic (montonic black *v*_7_-nullcline).

Before considering effects of reduced inhibition in the silent model, we note that once tonic inhibition is reduced, increasing it restores the normal breathing rhythm (Figure 9B). On the other hand, starting from the case of reduced tonic inhibition and introducing a small amount of recurrent inhibition keeps KF-s in an oscillatory state, and a further increase in *β*_7_ causes the *v*_7_-nullcline to become monotonic. Once this occurs, the KF-s system has a stable critical point at a *v*_7_ level such that *f*_7_(*v*_7_) *>* 0, corresponding to a tonic KF-s output (Figure 9B). Overall, in contrast to the tonic model, which suggests that inhibition within the KF is recurrent and hence depends on KF activity levels, the silent model suggests that inhibition to KF respiratory neurons is sustained and arrives from an outside source. Future experimental determination of the nature of inhibition to KF respiratory neurons will help to distinguish which, if any, of our models is consistent with the biological reality.

When the tonic inhibition, *b*_7_, to KF-s is reduced in the silent model, KF-s activity transitions from quiescent, with a stable critical point on the left branch of the *v*_7_-nullcline, to oscillatory, with an unstable critical point on the middle branch (Fig. 9A). The oscillations in KF-s provide oscillatory drive to post-I and produce a periodic breathing pattern similar to that observed in RTT. At the minimal value of *b*_7_ = 0.0, which we will refer to as our silent model representation of RTT, the apneic breathing pattern exhibited by the model is shown in Figure 10A. Notice that during RTT, the model does not produce late-E activation. This absence contrasts with the tonic model in Figure 2B, where we see late-E spiking during RTT-like apneas. The onset of periodic breathing, as *b*_7_ is reduced in the silent model, occurs at *b*_7_ ≈ 0.015 and the burst pattern just below this inhibition level is shown in Figure 10B. In Figure 10C,D we plot the period of KF oscillations, the apnea duration and the apnea frequency as the inhibition strength *b*_7_ is varied. The apnea duration remains roughly constant and is non-monotonic in *b*_7_ (shown in green Figure 10C), whereas the apnea frequency increases as *b*_7_ decreases (Figure 10D). These results agree qualitatively with the results for the tonic model (Figure 4C,D).

**Figure 10:**
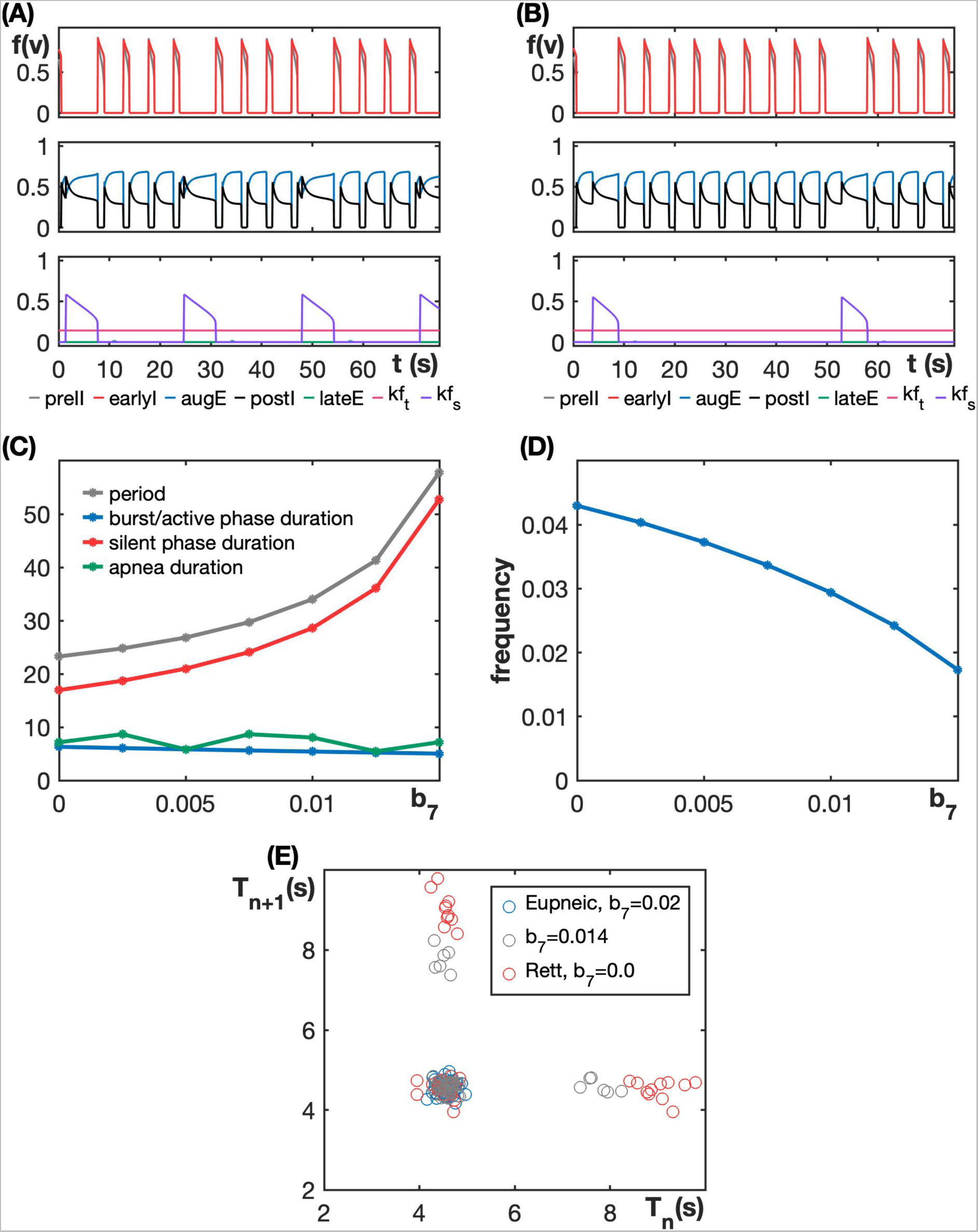
The silent model exhibits apneas, with preservation of normal cycle durations in between them, as tonic inhibition is reduced. (A) Periodic breathing exhibited by the model for *b*_7_ = 0.0 (RTT). (B) Respiratory pattern at an inhibition value near the point of transition of KF-s from tonic to oscillatory (*b*_7_ = 0.014). (C) The period of KF-s oscillations, KF-s burst/active phase duration, silent phase duration and apnea duration with respect to *b*_7_. (D) Apnea frequency, which is equal to the inverse of the period of KF-s oscillations, increases as the inhibition level to KF-s decreases.(E) Total respiratory duration during the (*n* + 1)*^st^* burst relative to that in the *n^th^* burst for silent model 1 with noise with *b*_7_ = 0.02 (default value), *b*_7_ = 0.014, and *b*_7_ = 0.0 (RTT).

Recall from Figure 2B and Figure 4A,B that as the inhibition level decreases in the tonic model, the KF-t burst/active phase duration decreases while the silent phase duration remains roughly the same. This causes the period of KF-t oscillations to decrease and apnea frequency to increase. In the silent model, however, we see in Figure 10A,B that the period of KF-s oscillations decreases mainly due to the reduction of silent phase duration of KF-s when *b*_7_ is lowered from 0.015. This effect arises because with less inhibition, the *v*_7_-nullcline in the silent phase moves farther away from the *m*_7_-nullcline (Fig. 9A), which yields less of a delay in the jump-up to the active phase. Therefore, this is another distinction between the tonic and silent models.

To study burst patterning in the silent model, we modified the model by adding Gaussian noise with mean 0 and standard deviation 1 by setting *s_i_* = 1 in (1) and (2). Figure 10E shows the total respiratory duration during the (*n* + 1)*^st^* burst relative to the duration for the *n^th^* burst (*T_n_*_+1_ versus *T_n_*) for inhibition levels *b*_7_ = 0.0 (RTT) and *b*_7_ = 0.014 (just after the onset of KF-s oscillations) and *b*_7_ = 0.02, the default value. The model did not show much variability in respiratory periods (blue dots in Figure 10E) during normal breathing conditions, as expected. Notice in Figure 10A that during KF-s oscillations, the inter-burst KF-s output matches the KF-s output level (≈ 0) during normal breathing. Therefore, in the RTT regime in this model, we expect that the breathing cycle durations in between apneas will remain the same as in the eupneic case. Aligning with this expectation, in the RTT case, we observe two different respiratory cycle durations (red dots in Figure 10E): apneas with duration ≈ 9s and normal breathing cycles during the silent phase of each KF-s oscillation, which match the cycle duration with the default model. This perseverance of the normal cycle duration even in the RTT condition differs from the activity in the tonic model since, in the tonic model under RTT conditions, the cycle duration in between apneas is shorter than the default respiratory breathing period.

For an intermediate value of inhibition in the silent model, for example *b*_7_ = 0.014, the apneas are shorter compared with the RTT case (Fig. 10E). We see in Fig. 10C, however, that apnea duration is clearly non-monotonic in *b*_7_. Notice from Fig. 10A,B that for different levels of *b*_7_, the initial KF-s activation can occur at different phases within the ongoing post-I active phase. For certain intervals of *b*_7_, this phase shift changes smoothly, but as *b*_7_ is decreased through other values, KF-s activation can switch from occurring early in every *n^th^* post-I cycle to late in every (*n* − 1)*^st^* cycle, yielding increased apnea durations. To avoid this arbitrary phase relation, we next introduced a putative excitatory connection from the post-I unit to the KF-s unit and examined its effects.

Including this excitatory drive does not change the eupneic breathing pattern exhibited by the silent model, since KF-s is silent in this regime (Figure 8). The effects of varying inhibition on the *v*_7_-nullcline and its intersection with the *m*_7_-nullcline also remain unchanged, and thus we focus on effects of varying tonic inhibition to KF-s. As was the case without the drive from post-I to the KF-s unit, when the inhibition strength parameter *b*_7_ is reduced sufficiently, the KF-s unit produces endogenous oscillations, which in turn leads to a perturbation in the normal breathing rhythm (Figure 11A). The transition of KF-s from the silent to the oscillatory state occurs now occurs at *b*_7_ ≈ 0.02 and the bursting pattern exhibited at this inhibitory level is shown in Figure 11B. Notice that in this case, post-I activation on certain cycles immediately recruits KF-s activation, without any phase lag. This effect smooths out the dependence of apnea duration on *b*_7_ and results in a monotonic relationship.

**Figure 11:**
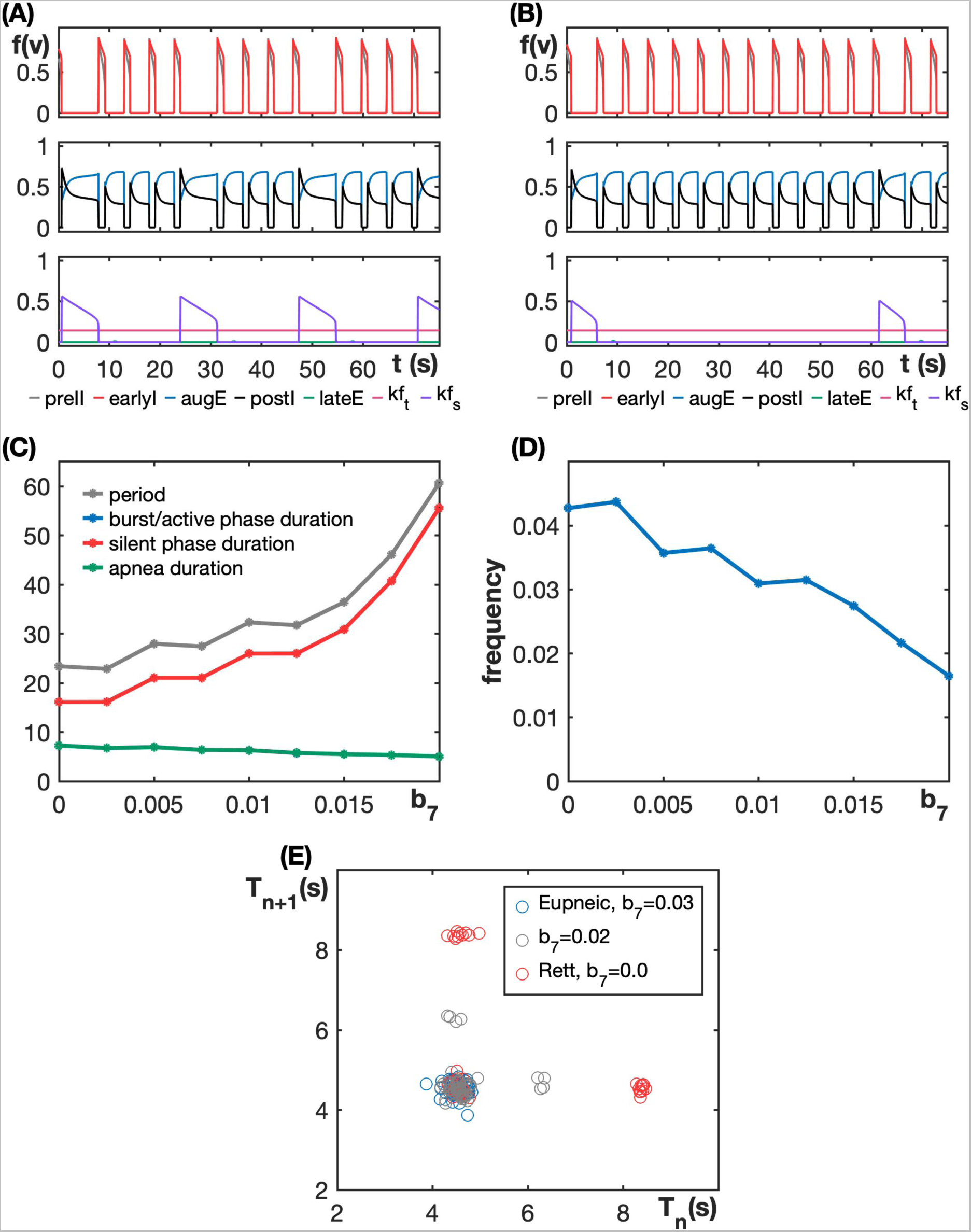
The inclusion of excitatory drive from post-I to KF-s reduces the variability in cycle durations. (A) Periodic breathing exhibited by the model for *b*_7_ = 0.0 (RTT). (B) Respiratory pattern at an inhibition value near the point of transition of KF-s from tonic to oscillatory (*b*_7_ = 0.02). (C) The period of KF-s oscillations, KF-s burst/active phase duration, KF-s silent phase duration, and apnea duration with respect to *b*_7_. In this case, the blue curve is identical to and completely hidden by the green curve, due to the connection from post-I to KF-s. (D) Apnea frequency increases non-monotonically as the inhibition level to KF-s decreases.(E) Total respiratory duration during the (*n* + 1)*^st^* burst relative to that in the *n^th^* burst when noise is included, with *b*_7_ = 0.03 (default value), *b*_7_ = 0.02 and *b*_7_ = 0.0 (RTT).

The results related to apnea duration and frequency (Figure 11C,D) qualitatively agree with those for the tonic model (Figure 4C,D) as well as for the silent model without feedback from post-I to KF-s (Figure 10C,D). The excitatory connection from post-I to KF causes the apnea duration to increase steadily as *b*_7_ is decreased in this augmented model (Figure 11C) and also results in a step-like decline in KF-s period as *b*_7_ decreases. This pattern represents a quantal effect: an approximately constant plateau occurs when KF-s activates once every *n* post-I cycles for fixed *n*, and then a step down to a lower period occurs when *b*_7_ becomes low enough that KF-s can activate once every *n* − 1 post-I cycles.

Figure 11E shows the burst patterning of the augmented silent model with Gaussian noise. This figure is comparable with Figure 10E in that the cycle durations between apneas carry over from the normal case to the cases with reduced inhibition levels. The new feature here is that the additional synaptic connection makes the cycle durations less variable than they were previously.

Finally, we do not separately consider the emergence of active expiration in the silent model. Since the KF-s population is silent by default, increasing the inhibition to KF-s from its baseline state would have no effect, while increasing the inhibition to KF-t with KF-s remaining silent would simply repeat the results that we obtained with the tonic model as described in Section 3.1.2.

## 4 Discussion

In this work, we consider a family of respiratory circuit models, which include the KF, designed to be minimal in terms of the complexity of the neural units, endogenous neuronal dynamics, and synaptic connections involved. We show that two of these models capture the central experimental findings on the response of respiratory patterns to manipulations involving the KF: (1) reduction of GABA-mediated inhibition to KF neurons induces periodic or intermittent breathing with apneas (Abdala et al., 2016; Dhingra et al., 2016); (2) durations of apnea events are maintained or lengthen with reductions in the inhibition level to KF (Abdala et al., 2016; Dhingra et al., 2016); (3) when intermittent breathing occurs, increases in KF inhibition can induce eupnea (Abdala et al., 2016, 2010); (4) from the eupneic state, additional increases in KF inhibition causes quantal activation of abdominal late-E activity (Koolen, 2021). Within this computational framework, we identify the synaptic connections that are compatible with the experimental results and obtain predictions about KF properties, circuit organization, and respiratory neuron behaviors under various conditions that could be tested to distinguish the two models and to check their validity. Our work builds on a series of assumptions and logical inferences, supported by the literature, which we now discuss before turning to the predictions that we derive as well as model limitations and future directions.

### 4.1 KF as a source of breathing irregularities

Intermittent breathing, such as observed in RTT, manifests as breathing that is periodically disrupted by apneic episodes. These apneas are accompanied by abnormally strong constrictor activity to the upper airways driven by motor neurons that exhibit a post-inspiratory activity pattern during regular breathing (Voituron et al., 2010;Abdala et al., 2010; Stettner et al., 2007). Post-inspiratory activity strongly depends on excitatory outputs from the KF (Smith et al., 2007; Dutschmann and Herbert, 2006). Disruption of inhibitory transmission in the KF, causing disinhibition, leads to periodic breathing with periods of excessive post-inspiratory activity to the larynx (Dhingra et al., 2016; Abdala et al., 2016). These findings provide strong (yet indirect) evidence that intermittent breathing can result from periodic KF overactivation on a time scale slower by an order of magnitude than respiration. Such a slow timescale suggests that the emerging oscillations can originate within the KF, resulting from a combination of intrinsic neuronal properties and synaptic inputs to the region. Our work illustrates some possible mechanisms that could give rise to this patterned KF activity.

### 4.2 Baseline activity in the KF

KF contains a large population of glutamatergic neurons that connects with the rCPG (Geerling et al., 2017). A recurrent excitatory network endowed with spike frequency adaptation properties (i.e., exhibiting a reduction in firing frequency within ongoing spiking activity) has the capacity to produce population-based rhythmic bursting. Changing parameters can transition such a system to a bursting regime regardless of whether it initially lies in a silent or tonically active state with a steady activity level. Therefore, we consider two basic KF network configurations. In the first configuration (*tonic model*), KF is represented by a single population with a tonic activity pattern under baseline conditions, which can transition into a bursting regime by relevant perturbations. In the second configuration (*silent model*), KF includes a subpopulation that is quiescent at baseline, either due to a a balance of endogenous currents that is inadequate to induce spiking without boosts in excitatory drive or due to suppression by ongoing inhibition, but starts producing oscillatory activity as parameters change. In the latter case, we assume that there is another population of neurons in the KF that always produces tonic output to provide the necessary excitatory drive to the rCPG. Still, unlike in the tonic model, this population does not become oscillatory under the manipulations that we consider.

### 4.3 Sources of GABA-mediated inhibition

Experiments show that reduced GABAergic inhibition in the KF leads to the emergence of slow oscillations modulating rCPG activity and causing breathing irregularities (Abdala et al., 2016). Depending on the KF network configuration, we infer different origins of this inhibition. Specifically, in the tonic model, this inhibition appears to be recurrent. Although we implement this recurrence as self-inhibition in our minimal model, this recurrent inhibition is a reduced representation of a scenario in which the KF population excites inhibitory interneurons that in return provide feedback inhibition. The location of these inhibitory interneurons is not specified by our model. One can speculate that these neurons can represent a local inhibitory subpopulation providing negative feedback within the KF for self-regulation. Alternatively, they could be located in another site, such as the BötC or the parabrachial nucleus, proved to have recurrent connectivity with KF (Ezure et al., 2003; Gaytán et al., 1997; Varga et al., 2020). In contrast, in the framework of the silent model, the GABAergic inhibition in KF is predicted to take the form of a tonic or sustained drive, probably originating externally to the KF; for example, the NTS is known to send feedforward inhibitory projections to the KF and hence could represent the source of this input (Otake et al., 1992; Kubin et al., 2006).

### 4.4 Mechanisms of 5-HT**_1_**_A_ inhibition

Experimental evidence indicates that manipulating 5-HT_1A_ receptors in the KF produces changes in respiratory activity comparable to the changes induced by modulating GABAergic inhibition (Dhingra et al., 2016; Abdala et al., 2016). Specifically, 5-HT_1A_ antagonists evoke intermittent breathing while 5-HT_1A_ agonists can reduce breathing irregularities or even restore eupnea (Dhingra et al., 2016; Besnard et al., 2012). While serotonergic inputs originate outside of the KF (e.g., in the raphe nucleus), their functional effects, mediated by 5-HT_1A_ receptors, are compatible with both the recurrent and tonic inhibition scenarios. Specifically, tonic inhibition could be membrane hyperpolarization due to the activation of 5-HT_1A_R-coupled potassium channels (Montalbano et al., 2015), which is compatible with the silent model. On the other hand, another known impact of 5-HT_1A_R activation is the enhancement of glycinergic inhibition in neurons that express the GlyR *α*3 subunit (Shevtsova et al., 2011). This mechanism could be involved in altering the recurrent inhibition provided by glycinergic interneurons. Potentially, the interneuron population could coexpress both GABA and glycine and thus could mediate the effects of both GABAergic and serotonin-modulated recurrent inhibition in the tonic model. Finally, our results on the emergence of active expiration (Section 3.1.2) suggest that 5-HT_1A_ agonists may strengthen the recurrent inhibition within a tonic population of KF neurons.

### 4.5 Additional predictions and implications

Both the tonic and silent models, with appropriate forms of inhibitory connections, capture the set of benchmarks that we initially imposed. While it is not feasible to use these models for quantitative predictions, our modeling framework can allow us to propose experimental designs that can be deployed to test and select between these models. For starters, the structural differences that distinguish the two models themselves represent predictions. Specifically, the tonic model assumes that the KF would feature prominent recurrent inhibition, through which increased activity in KF neurons strengthens the inhibitory inputs that these neurons receive. In contrast, the silent model predicts an important role for a sustained inhibitory input to KF from an outside source, as well as the presence of a subpopulation of KF neurons that exhibit little or no activity under control states of eupneic respiration.

The network configurations predicted by the two models also have different implications for abdominal activity during periods of intermittent breathing when inhibition in KF is compromised. Indeed, in the tonic model, reduction in recurrent inhibition transforms steady KF activity into an oscillatory regime with periods of overactivity inducing temporary apnea, adaptation to near a normal activity level, and then periods of silence (see Fig. 4A, B). During the latter periods, KF does not provide excitation to post-inspiratory neurons in BötC, which results in a disinhibition of expiratory neurons in pFL and evokes abdominal late-E activity that drives active expiration (Fig. 4A, B). In contrast, in the silent model, oscillations in KF emerge in a previously silent population while the steady activity of the tonic KF subpopulation remains unaltered. Therefore, conditions for the abdominal activity breakthrough are never created (see Fig. 10A, B). Interestingly, some apneic events in some RTT mice are accompanied by abdominal activation without a clearly corresponding late-E activation (Abdala et al., 2010), but this result does not suffice to distinguish the two models, as late-E activity could be present yet phase-shifted by other factors.

Typically, experiments relating to respiration in RTT and under compromised KF inhibition compare across two groups, such as control versus experimental mice or wild-type versus knock-out mice. Using computational models allows us to generate predictions about the effects that we expect from gradual changes that induce states in-between the extreme endpoints. In the tonic model, reductions in inhibition to KF induce a regime of KF oscillations with a long epoch of sustained KF activity within each cycle. In this regime, apneas are interspersed with respiratory cycles with a mix of durations, including some slightly longer and some slightly shorter than those seen under control conditions (Figure 4). Further decreases in inhibition are predicted to shorten the KF active duration and the oscillation period, with a small increase in apnea duration and a regularization of cycle durations in between apneas. The silent model points to a different pattern of changes with progressive decrease in inhibition to KF. In this model, we see that when the onset of apneas occurs, the durations of the cycles in between the apneas are fairly stereotyped and match those seen in control conditions (Figure 10). Decreasing inhibition allows apnea durations to show a net increase, but unlike in the tonic model, this relationship is non-monotonic. If we hypothesize the inclusion of an additional excitatory connection from BötC neurons to the quiescent KF population, then we recover a more consistent phase relation between expiratory BötC and KF activity, which regularizes the trend in apnea durations (Figure 11).

In summary, the tonic and silent models are both consistent with published experiments but can be differentiated in terms of their predictions about KF activity in baseline conditions, inhibitory pathways that impact KF activity, and alterations in respiratory patterns resulting from decreases as well as increases in inhibition to the KF. Finally, an interesting possibility that should also be kept in mind is that the tonic and silent models may both be valid but in different regimes of respiratory circuit function; for example, changes under vagotomy may alter KF activity in a way that corresponds to switching between these two models. These possibilities require experimental verification to be validated.

### 4.6 Model limitations and future directions

By starting from a minimal modeling framework, we were able to use analytical tools, including analysis of nullclines in certain phase planes, to argue against the presence of specific combinations of KF endogenous dynamics and forms of inhibition to the KF, while also providing a proof of principle that certain other configurations can capture a range of experimental findings and merit further consideration. A natural next computational step would be to address the limitations of this minimal modeling framework by implementing the proposed circuit arrangements in a more complete model featuring populations of spiking neurons with a full complement of known transmembrane ion currents in each of the constituent brain regions as well as more biologically detailed synaptic interactions. Past work has shown that endowing KF neurons with intrinsic bursting capabilities and also assuming the presence of certain synaptic interactions between neurons in the KF, in the parabrachial nucleus (PBN), and in the BötC produces a circuit that also produces eupneic respiratory output and captures the effects of vagotomy as well as periodic breathing following reduction of GABAergic inputs to the KF (Wittman et al., 2019). Because these ideas were considered previously and represent a specific set of assumptions that go beyond our minimal framework (e.g., involvement of a PBN component), we did not consider these properties in this work. While the results that we obtained show that endogeneous KF oscillations are not necessary to explain experimental findings, since previous studies have demonstrated the presence of KF neutrons with phasic discharge patterns both with and without vagotomy (Mörschel and Dutschmann, 2009), another natural future direction will be to compare these frameworks directly and to explore possible ways to integrate these models. Of course, future experimental work to directly derive more information about the intrinsic properties of KF neurons, their endogenous dynamics, and the synaptic interactions in which they are involved would be an invaluable complement to computational approaches. Hopefully the findings from this work and other computational studies can guide these experimental investigations in productive directions.

### 4.7 Conclusions

Our modeling results, tuned to capture experimental observations, highlight the importance of the KF neurons in maintaining eupneic breathing and support the hypothesis that disruptions in KF activity can contribute to the emergence of pathological respiratory phenotypes. Our predictions suggest two different possible circuit organizations that may be present within the KF, with neuronal subpopulations displaying distinct intrinsic characteristics and synaptic connections. These new possibilities can foster future experimental and pre-clinical studies examining in detail the proposed features and their involvement in pathological states associated with breathing irregularities, such as RTT.

## 5 Additional Information

### 5.2 Data Availability

All the code and simulation results will be made available upon request.

### 5.3 Author contributions

S.J. and W.B. generated and analyzed data and drafted/revised the work; D.Z. and A.A. contributed to the conception of the work and revised the manuscript critically for important intellectual content; J.R. and Y.M. conceptualized and designed the work, analyzed and interpreted data and drafted the work. All authors approved the final version of the manuscript. All authors agree to be accountable for all aspects of the work in ensuring that questions related to the accuracy or integrity of any part of the work are appropriately investigated and resolved. All persons designated as authors qualify for authorship, and all those who qualify for authorship are listed.

